# Effect of feeding and thermal stress on photosynthesis, respiration and the carbon budget of the scleractinian coral *Pocillopora damicornis*

**DOI:** 10.1101/378059

**Authors:** Niclas Heidelberg Lyndby, Jacob Boiesen Holm, Daniel Wangpraseurt, Renaud Grover, Cécile Rottier, Michael Kühl, Christine Ferrier-Pagès

## Abstract

Studying carbon dynamics in the coral holobiont provides essential knowledge of nutritional strategies and is thus central to understanding coral ecophysiology. In this study, the first aim was to investigate the effect of daily feeding and thermal stress on oxygen (O2) rates measured at polyp-scale with microsensors and at whole fragment scale using incubation methods. The second aim was to assess the carbon budget of the symbiotic association using H^13^CO_3_, under the different conditions. Micro- and macro-scale measurements revealed enhanced O_2_ evolution rates for fed compared to unfed corals. However, gross O_2_ production in fed corals was increased at high temperature on a macroscale but not on a microscale basis, likely due to a heterogeneous distribution of photosynthesis over the coral surface. Starved corals always exhibited reduced photosynthetic activity at high temperature, which suggests that the nutritional status of the coral host is a key limiting factor for coral productivity under thermal stress. Quantification of photosynthate translocation and carbon budgets showed very low incorporation rates, for both symbionts and host (0.03 - 0.6 μg C cm^-2^ h^-1^) equivalent to only 0.008 - 0.6 %, of the photosynthetically fixed carbon for *P. damicornis*, in all treatments. Carbon loss (via respiration and/or mucus release) was about 41 - 47 % and 52 - 76% of the fixed carbon for starved and fed corals, respectively. Such high loss of translocated carbon suggests that *P. damicornis* is nitrogen and/or phosphorus limited. Heterotrophy might thus cover a larger portion of the nutritional demand for *P. damicornis* than previously assumed. Our results suggest that active feeding plays a fundamental role in metabolic dynamics and bleaching susceptibility of corals.

## Introduction

Coral reefs in tropical to subtropical waters are primarily founded by calcifying (scleractinian) corals that are fueled by an endosymbiotic relationship between the cnidarian host and their photosynthesizing dinoflagellates, commonly referred to as zooxanthellae, belonging to the genus *Symbiodinium*. This mutual association enables corals to thrive in oligotrophic waters by translocation of photosynthates from the symbiont to the animal host (Muscatine *et al*., 1981; Tremblay *et al*., 2012). In return, the zooxanthellae receive shelter and inorganic nutrients via waste products from the coral host metabolism (Muscatine & Cernichiari, 1969).

It is generally accepted that symbiont photosynthesis covers up to 95% of the daily energy demand of the animal host (Muscatine *et al*., 1984), especially in shallow waters where photosynthetically active radiation (PAR) is not limiting. However, corals that inhabit deeper and/or turbid waters, are far more dependent on active feeding, as photosynthesis becomes light limited (Falkowski *et al*., 1984; Palardy *et al*., 2008). Studies have shown that corals utilize heterotrophy for supplementary carbon and nutrient acquisition, even in light saturated environments, thus implying an advantageous effect from active feeding (Yonge & Nicholls, 1931; Wellington, 1982; Sebens *et al*., 1996; Houlbrèque *et al*., 2004). Furthermore, the animal host relies on heterotrophy for obtaining nitrogen, phosphorus and other essential nutrients, that are not acquired through autotrophy (Houlbrèque & Ferrier-Pagès, 2009).

Heterotrophy becomes particularly important during bleaching events, as corals rely on zooxanthellae to sustain their daily metabolic energy demand and the loss of symbionts induce coral starvation and can subsequently lead to coral death (Brown, 1997). In addition, coral bleaching can lead to an ‘optical feedback loop’ (Enríquez *et al*., 2005), where symbiont loss promotes increased tissue light penetration and skeletal backscattering, which subsequently leads to enhanced light absorption by the remaining zooxanthellae (Enríquez *et al*., 2005; Wangpraseurt *et al*., 2017). This can lead to additional photodamage and production of reactive oxygen species (ROS), which can further accelerate the bleaching process (Weis 2008; Lesser 1996). In such conditions, heterotrophy increases resilience to bleaching by storage of long lasting energy deposits (Hughes & Grottoli, 2013), and sustains basic coral host metabolism and the recruitment and re-colonization of zooxanthellae when more favorable conditions are present (Grottoli *et al*., 2006; Rodrigues & Grottoli, 2007; Hughes *et al*., 2010). Therefore, investigations of the proportional input from heterotrophy compared to autotrophy during coral bleaching reveal a potentially very high (up to 100%) contribution of heterotrophically acquired carbon to daily animal respiration (CHAR) in certain species (Grottoli *et al*., 2006; Courtial *et al*., 2017). This implies the existence of species-specific adaptions among corals to utilize active feeding as a primary nutrient source under stressed conditions.

Many other factors interact with heterotrophy to mitigate coral bleaching. For example, corals can receive an enhanced transfer of photosynthates by the remaining symbionts (Tremblay *et al*., 2016), or decrease their metabolism (as assessed by the decrease in respiration rates and carbon release) during stress. These parameters can be assessed by making the carbon budget of corals, through the use of ^13^C-isotope tracer experiments (Tremblay *et al*., 2012), and O_2_ fluxes. The combination of the two techniques allows the quantification of the autotrophic carbon fixation rates, of the fraction stored in either symbionts or host tissue, or lost through respiration and mucus release, and yield valuable insight on the current health and productivity state of the coral holobiont. One limiting step in these fluxes is the right determination of the O_2_ fluxes, or in other word, the right determination of the respiration and gross photosynthesis rates. Most measurements indeed tend to focus on a single spatial scale (e.g. whole colony scale logging), and assume a homogenous distribution across the surface. However, O_2_ produced by symbiont photosynthesis is often used immediately for symbiont and coral respiration. This intimate association between autotrophic and heterotrophic processes makes the quantitative and spatial separation of respiration and photosynthesis difficult (Kühl *et al*., 1995, 1996). A method for measuring gross photosynthesis independent of respiration is available based on the use of O_2_ microelectrodes (Revsbech & Jørgensen, 1983). In addition, gross photosynthesis is dependent on the light reaching the symbionts (Wangpraseurt *et al*., 2015), but the fate and turnover rate of the incident irradiance reaching the coral surface (radiative energy budget), is still poorly known. Establishing a radiative energy budget allows the determination of the energy lost to the system by reflection or absorbed in the coral tissue where it can either be dissipated as heat or used for photosynthesis (Brodersen *et al*., 2014; Lichtenberg, Brodersen & Kühl, 2017; Lyndby *et al*., submitted).

In this present study, the main aim is to investigate O_2_ rates at different spatial scales by employing a combination of microsensor and incubation techniques to examine the effect of daily feeding and thermal stress on coral bleaching susceptibility. Furthermore, we establish the first carbon and radiative energy budgets on *P. damicornis*, a frequently used model species in coral ecophysiology. This part of the study will focus on the implication of thermal stress and feeding on the carbon budget, whereas an accompanying paper (Lyndby *et al*., submitted) will focus on the radiative energy budget of *P. damicornis* under same conditions.

## Materials and methods

*Pocillopora damicornis* corals cultured at the Centre Scientifique de Monaco (CSM), were prepared by cutting eight mother colonies into ^~^120 fragments of 2-3 cm in diameter, and hung from nylon threads in the tanks. All corals were kept under white light (250W metal halide lamps), at a downwelling irradiance (400-700 nm) of 200-250 μmol photons m^-2^ s^-1^, illuminated for 12 hours a day. Eight tanks (water renewal rate of 10 L h^-1^, 25°C, salinity of 35) were set up two months in advance of the bleaching experiment as to randomly divide the corals into two groups of four tanks each with either fed or unfed corals. Fed corals were fed once a day with ^~^4000 *Artemia* nauplii per coral fragment, four times per week, and unfed corals were kept unfed throughout the study, including the two months prior to measurements.

About 20 fragments were kept in each control tank, while three thermal stress tanks contained a total of 40 fragments for each feeding treatment. At the beginning of the study, all eight tanks started at 25°C. The tanks used for thermal stress were then gradually ramped up to 30°C over a period of five days, increasing 1°C per day. Thermally stressed corals were kept at 30°C for an additional two to three days before starting measurements. Corals measured on after the two-three days of 30°C thermal stress are denoted as time point 1 (T_1_ shortened). After an additional five days of thermal stress, measurements were conducted again, denoted as time point 2 (T_2_). Measurements on control corals were done in between the waiting period for T_1_ and T_2_, and are denoted time point 0 (T_0_).

### Experimental setup and approach

Corals were placed in a black acrylic flow chamber for all microsensor measurements. The flow chamber was supplied with seawater (25°C, salinity 35) from a heated water reservoir (10 L) at a flow rate of ^~^0.25 cm s^-1^ through the chamber. A motorized micromanipulator (MU-1, PyroScience, Germany) was attached to a heavy-duty stand to facilitate positioning of microsensors on fragments in a 45° angle relative to the vertically incident light. A digital microscope (Dino-Lite Edge AM7515MZTL, AnMo Electronics Corporation, Taiwan) was used to carefully position the sensor tip on the coral tissue surface (See figure 1e-f in accompanying paper by Lyndby *et al*., submitted). A tungsten halogen lamp (KL-2500 LCD, Schott, Germany), fitted with a fiber light guide and collimating lens, was used to illuminate the flow chamber with white light (See Supplementary Figure 1) vertically from above. A calibrated spectroradiometer (MSC15, GigaHertz-Optik, Germany) was used to quantify the absolute downwelling photon irradiance at different lamp settings (80, 167, 250, 480, 970, and 2400 μmol photons m^-2^ s^-1^), and in addition storing the radiometric energy spectra in W m^-2^ nm^-1^. The intensity was adjusted via the aperture of the lamp without spectral distortion. The setups were covered with black cloth during measurements. For each coral used in this study, microsensor measurements were done on three randomly chosen polyps located on the branch tips (See dotted circles in figure 1a-b in accompanying paper by Lyndby *et al*., submitted).

**Figure 1.**
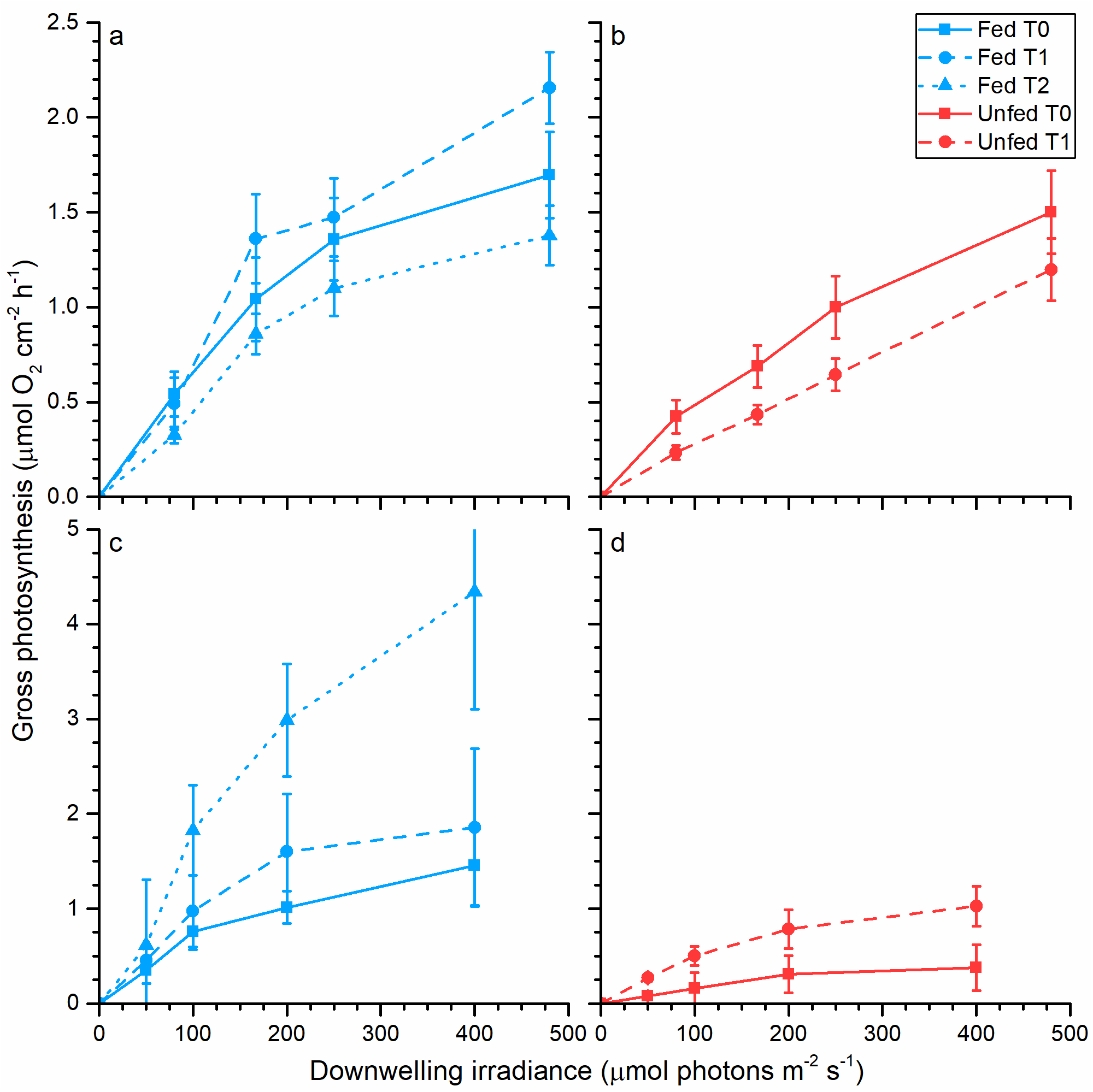
Gross O_2_ production in *Pocillopora damicornis* as a function of increasing downwelling photon irradiance as measured with O_2_ microsensors on a polyp scale (a-b) and with O_2_ gas exchange measurements of whole coral fragments in incubation chambers (c-d). Measurements were performed on daily fed corals (a, c) and unfed corals (b, d) for each of three temperature treatments; T_0_ (control, solid lines), T_1_ (3 days at 30°C; dashed lines) and T_2_ (8 days at 30°C; dotted lines). Symbols with error bars represent mean ± SEM (n = 5-12).

### Microsensor measurements

Gross photosynthesis was measured with Clark-type O_2_ microelectrodes with a tip diameter of ^~^25 μm, a low stirring sensitivity (<5%), and a fast response time (<0.5 s; OX-25 fast, Unisense, Denmark). The microelectrodes were connected to a pA meter (Unisense, Denmark), and were linearly calibrated from readings in 100% air saturated seawater and anoxic water (by addition of Na2SO3) at experimental temperature. The percentage of O_2_ air saturation was converted to μmol L^-1^ using gas tables for the O_2_ solubility in air-saturated water as a function of temperature and salinity (www.unisense.dk). For gross photosynthesis estimates, data were recorded on a strip-chart recorder (BD25, Kipp&Zonen, Netherlands) connected to the picoammeter. Gross photosynthesis was estimated using the light-dark shift method as described in detail by Revsbech & Jørgensen (1983). O_2_ measurements performed at the coral tissue surface were regarded as representative of the entire tissue volume of *P. damicornis* given that the average tissue was ^~^200 μm thick, which approximates the spatial resolution of the light-dark shift method during the 2-3 second period of darkening.

For measurements of holobiont dark respiration, coral fragments were dark adapted for 15 min before measuring O_2_ concentration profiles across the diffusive boundary layer (DBL) between the mixed overlaying water and the coral tissue surface. Measurements started at the tissue surface (depth = 0 μm) and the microsensor tip was moved upwards in steps of 50 μm (10 s rest time per measuring position) until reaching the ambient water with a constant O_2_ concentration.

Subsequently, net photosynthesis vs. irradiance curves were measured under 7 increasing levels of incident photon irradiance (E_d_(PAR) = 0, 80, 167, 250, 480, 970, and 2400 μmol photons m^-2^ s^-1^). Each coral fragment was incubated for 10 min at each irradiance level, ensuring steady-state conditions. A total of 4 replicate fragments were measured per treatment and sample times using 3 randomly selected coenosarc and polyp tissue areas, i.e. a total of 24 measurements per treatment. Towards the end of the bleaching treatment, some fragments showed no O_2_ production on the coenosarc and we thus limited the replicate measurements for those fragments.

Local net-photosynthesis and dark-respiration rates were determined as the net O_2_ flux, *J* (nmol O_2_ cm^-2^ s^-1^), as calculated from the measured steady-state O_2_ profiles using Fick’s first law of diffusion:

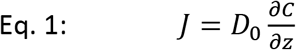

where *D*_0_ is the molecular diffusion coefficient of O_2_ in seawater at experimental salinity and temperature (*D*_0_ = 2.2327 · 10^-5^ cm^2^ s^-1^ at 25°C and S=35, and *D*_0_ = 2.5521 · 10^-5^ cm^2^ s^-1^ at 30°C and S=35; Broecker & Peng, 1974), δC/δz is the slope of the linear part of the O_2_ concentration gradient in the DBL, defined by the change in O_2_ concentration (δC) over a specific depth interval (δz).

### Gas exchange measurements

Additional measurements of respiration and photosynthesis were made by measuring the net O_2_ gas exchange of coral fragments (Hoogenboom *et al*., 2010) under photon irradiance levels of 0, 50, 100, 200 and 400 μmol photons m^-2^ s^-1^. For this, coral fragments (3-5 for each treatment) were positioned upright in small incubation chambers (50 mL) and the fragments were illuminated from the side, with a 250W metal halide lamp. To ensure even illumination of the entire coral fragment, aluminium foil was placed behind the incubation chamber, reflecting the incident beam and illuminating the otherwise shaded areas of the coral fragment. The photon scalar irradiance in the incubation chamber was measured for defined lamp settings using a spherical mini quantum sensor (sphere diameter of 3.7 mm, US-SQS/L, Walz GmbH, Germany) attached to a light meter (LI-190, Li-Cor, Lincoln, NE, USA).

The temperature in the experimental chambers was kept at 25°C for control and at 30°C for heat-treated fragments and the chambers were constantly stirred with a magnet. The O_2_ concentration was logged over time with an O_2_-optode (O_2_ MiniOptode, Unisense A/S, Aarhus, Denmark) connected to a computer with Oxy-4 software (4-channel fiber-optic O_2_ meter, PreSens, Regensburg, Germany). O_2_ optodes were linearly calibrated from measurements in air saturated water (bubbled with air) and anoxic water (flushed with N_2_) according to the manufacturers recommendations.

Dark holobiont respiration was determined as the O_2_ depletion after 15 min dark adaptation and light respiration as the initial post-illumination O_2_ depletion after each experimental irradiance level. Net photosynthesis (*P_n_*) was determined as the O_2_ concentration increase after 15 min illumination at the given photon irradiance similar to light respiration measurements. The O_2_ depletion and production curves were fitted with linear regression and rates of respiration and net-photosynthesis were calculated as:

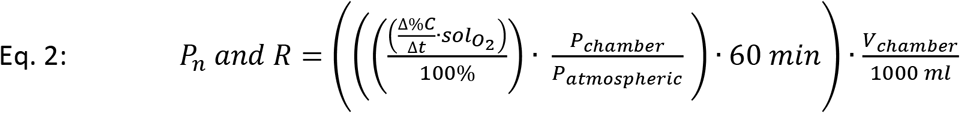

Δ%*C*/Δ*t* is the change in % O_2_ over time (% min^-1^), *sol*_*O*_2__ is the O_2_ solubility (μmol L^-1^) for the given temperature and salinity ( *sol*_*O*_2__ = 211.3 μmol/L at 25°C and S=35, and *sol*_*O*_2__ = 194.6 μmol/L at 30°C and S=35), *P*_chamber_ / *P*_atmospheric_ is the relative difference in partial pressure (hPa) between atmospheric air and the water mass in the incubation chamber, *V*_Chamber_ is the volume of the chamber (mL), and *P_n_* and *R* are rates of net-O_2_ production and O_2_ consumption, respectively (μmol O_2_ h^-1^). Gross photosynthetic (*P_g_*) rates (in μmol O_2_ h^-1^) were then estimated by adding *R* to *P_n_*.

Besides gas exchange measurements on intact coral fragments, we also measured O_2_ gas exchange on freshly isolated symbionts from 3 previously measured fragments. The symbionts were extracted from 3 fragments per treatment and time point, by first removing the coral tissue using an airbrush with 35 mL filtered seawater (FSW; 0.45 μm). Subsequent homogenization of tissue with a Potter tissue grinder, and transfer of the tissue slurry to a 50 mL Falcon^®^ tube that was centrifuged for 10 min at 850*g*. After removal of the supernatant the remaining symbiont pellet was resuspended in FSW (0.45 μm).

The symbiont density was quantified in a 100 μL sub-sample using a Z1 Coulter Particle Counter (Beckman Coulter). The O_2_ fluxes were estimated using similar techniques as whole fragment measurements described previously, and were converted to carbon units according to Anthony & Fabricius (2000).

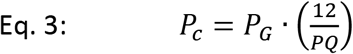

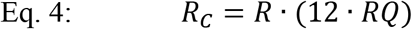

Where *P_c_* is the total amount of inorganic C fixed through photosynthesis, *P_G_* is the O_2_ gross production calculated from *P_n_* and *R* in μmol O_2_ cm^-2^ h^-1^; 12 is the mass of a carbon atom, *PQ* is the photosynthetic quotient, here assumed to be 1.1 mol O_2_:mol C (Muscatine *et al*., 1981), *R_C_* is the rate of consumed inorganic C (μg C m^-2^ h^-1^) by respiration *R*, and *RQ* is the respiratory quotient, here assumed - equal to 0.8 mol C:mol O_2_ (Muscatine *et al*., 1981).

### H^13^CO_3_ tracer incubations

Corals used for stable isotope incubations with H^13^CO_3_ were mounted in beakers incubated with FSW (controls) or ^13^C-enriched FSW (treatments) for pulse periods of 4 hours (Tremblay *et al*., 2012). The treated fragments were exposed to a solution of 0.6 mM NaH^13^CO_3_ (98% ^13^C; #372382, Sigma-Aldrich, St Louis, MO, USA), mixed with 200 mL FSW (Tremblay *et al*., 2012). The beakers were left open and the solutions were constantly stirred by a magnet bar to avoid re-use of respired ^13^C.

The beakers with fragments of *P. damicornis* (3-5 for each treatment) were kept at a constant temperature of 25°C for control and 30°C for heat-stressed corals by a water bath under thermal regulation, and were illuminated with a constant incident photon irradiance of 200 μmol photons m^-2^ s^-1^. During these incubations, O_2_ production and consumption were not recorded, but the rates of carbon fixation and respiration were derived from the above mentioned O_2_ gas exchange measurements at 200 μmol photons m^-2^ s^-1^. For quantification of ^13^C enrichment in coral fragments, the tissue was detached from the skeleton by an air brush with 10 mL FSW. The tissue slurry was homogenized using a Potter tissue grinder and the animal and symbiont fractions were then separated by centrifugation (1328g for 5 min). To avoid tissue contamination, the symbionts were rinsed several times in FSW and checked under a microscope. Tissue and symbionts were separated, frozen in liquid N2 and freeze dried for later analysis. The ^13^C enrichment and carbon content in the animal tissue and symbionts were quantified with a mass spectrometer (Delta Plus, Thermo Fisher Scientific, Bremen, Germany) coupled via a type III interface with a C/N analyzer (Flash EA, Thermo Fisher Scientific), and compared with ^13^C/^12^C in the control fragments (Tremblay *et al*., 2012).

### Carbon budget calculations

Incorporation rates in symbionts and host (ρ; defined as the rate of incorporated C per area and time, i.e., μg C cm^-2^ h^-1^) were calculated according to Tremblay *et al*. (2012), by the following equations:

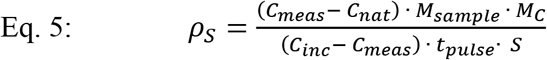

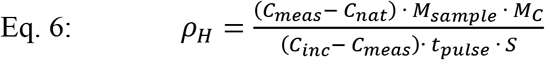

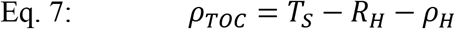

where *ρ* refers to the carbon incorporation rate (μg C cm^-2^ h^-1^) into the symbionts (*ρ_S_*, Eq. 5) and coral host (*ρ_H_*, Eq. 6), and *ρ_TOC_* is calculated as the released C from the host (μg C cm^-2^ h^-1^).

*T_S_* is the translocation rate from symbiont to host, *C*_meas_ and *C*_nat_ are the percentage of ^13^C measured in our experimentally treated samples (symbionts and host tissue) and the control samples, respectively, *M*_sample_ is the mass of the freeze-dried sample (mg), *M*_C_ is the mass of carbon per milligram of symbiont or host tissue (μg mg^-1^) or the release of C from the coral holobiont (μg). *S* denotes the surface area (in cm^2^) of the coral fragment as quantified by the wax dipping technique (Veal *et al*., 2010), tpulse is the incubation time (hours) of the fragments in control and enriched incubation medium, and *C*_inc_ is the percent ^13^C of the DIC in the medium. We used seawater that contained 2 μM HCO^-^_3_ with a natural ^13^C abundance of 1.1285% and added 0.6 μM HCO_3_^-^ with a ^13^C abundance of 98%. *C*_inc_ was then determined by the following equation:

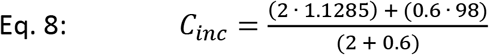

Based on the previous determined incorporation rates, respiration rates and the amount of fixed inorganic C going into the symbionts, the amount of C translocation from the symbionts to the coral host was calculated as the net remaining C after symbiont respiration and incorporation:

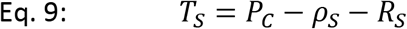

See Figure 3 for schematic overview of the carbon budget and its pathways.

### Statistical analysis

Photosynthetic rates and carbon fluxes were tested for homogeneity of variance using a two-way *F*-test and for significant differences using a two sample Student’s *t*-test (for equal variances) or Welch’s *t*-test (for unequal variances). Statistical tests were performed in OriginPro 9.1 (Origin, USA).

## Results

### Photosynthesis vs. irradiance curves

O_2_ production was enhanced in fed *vs* unfed *Pocillopora damicornis* corals for polyp-scale microsensor (microscale) and whole fragment incubation (macroscale) measurements (Figure1). Fed corals from time point T_0_ and T_1_ (Figure 1a, c) exhibited higher gross photosynthesis for both measuring techniques with small variations (± 0.1-0.4 μmol O_2_ cm^-2^ h^-1^) compared to unfed corals that showed a more profound variation between the techniques (± 0.5-0.8 μmol O_2_ cm^-2^ h^-1^; Figure 1b, d). The effect of thermal ramping on fed corals led to a 20% increase in O_2_ production from T_0_ to T_1_. However, on a longer term (T_2_), we found a significant effect of increased temperature at the macroscale, leading to a 2.2-fold enhancement in O_2_ production at T_2_ (4.1 ± 0.75 vs 1.3 ± 0.2 μmol O_2_ cm^-2^ h^-1^) as compared to T_0_ (Student’s *t*-test: *p* < 0.001; Figure 1c). The O_2_ production at microscale was on the contrary reduced by 0.6 ± 0.3 μmol O_2_ cm^-2^ h^-1^ from T_1_ to T_2_ (Figure 1a), which deviates from the observed gross photosynthesis on a macroscale at T_2_ (Figure 1a, c). O_2_ production was reduced for unfed corals on both micro- and macroscale.

Gas-exchange measurements of areal net photosynthesis showed onset of net O_2_ production in fed corals at a photon irradiance >100 μmol photons m^-2^ s^-1^ for both micro- and macroscale measurements (Figure 2a, c). In contrast to fed corals, unfed corals only exhibited net O_2_ production at irradiances >150 μmol photons m^-2^ s^-1^ (Figure 2b, d). The combination of temperature increase and starvation led to a slightly higher compensation irradiance (i.e. the irradiance above which net O_2_ production is observed) at 250 μmol photons m^-2^ s^-1^ at T_0_, and 400 μmol photons m^-2^ s^-1^ at T_1_ on a microscale, and 150 μmol photons m^-2^ s^-1^ at T_0_-T_1_ on a macroscale (Figure 2b, d). We found enhanced light respiration with microscale measurements for both fed and unfed corals (Figure 2a, b), as compared with macroscale measurements. Dark holobiont respiration was similar for all treatments except for fed corals at T_2_ (Figure 2c), which showed a 3-fold increase in O_2_ consumption compared to fed corals at T_0_ (0.6 ± 0.05 vs 2.4 ± 0.25 μmol O_2_ cm^-2^ h^-1^).

**Figure 2.**
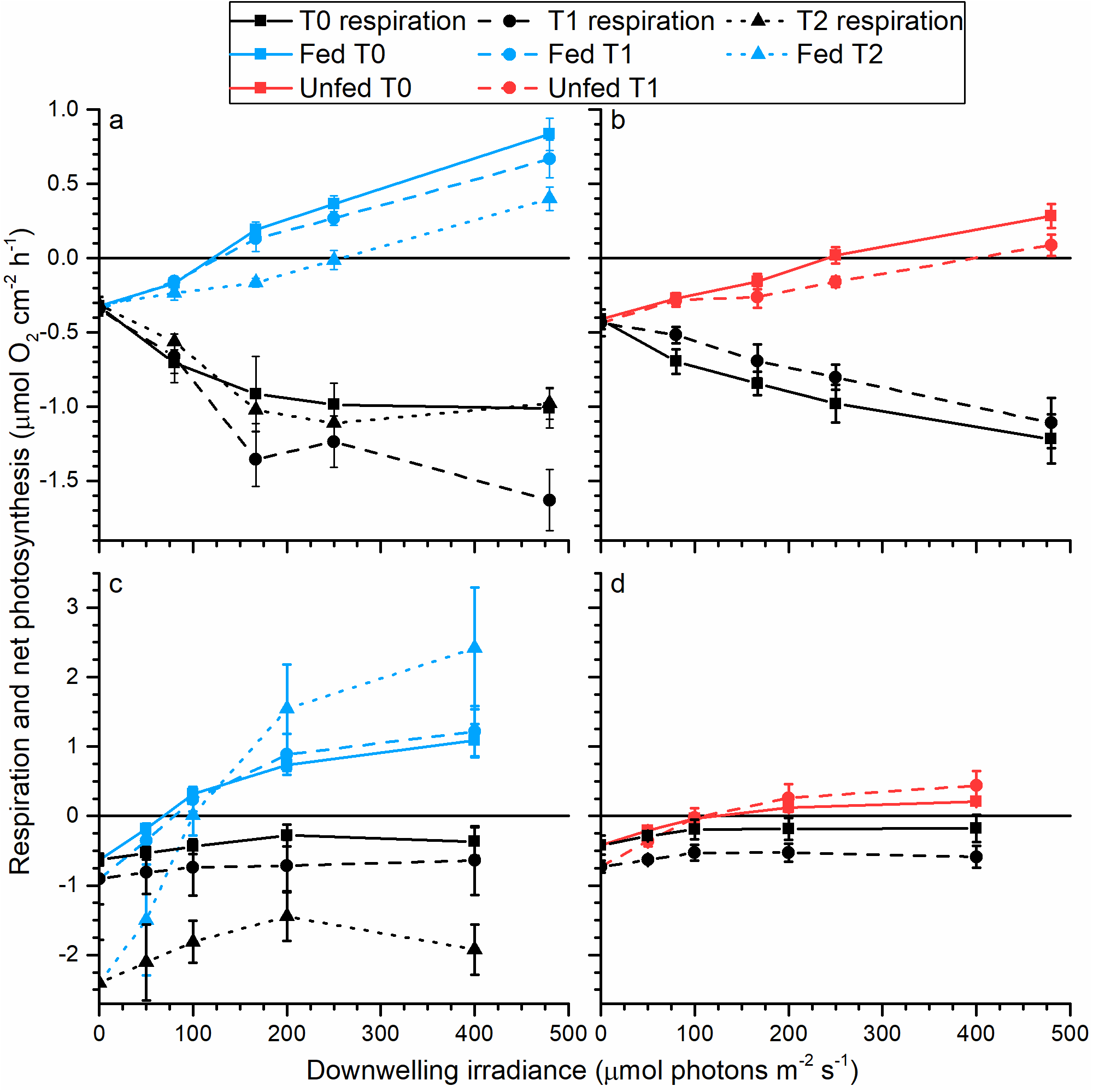
Net O_2_ production and consumption in *Pocillopora damicornis* as a function of increasing downwelling photon irradiance measured with O_2_ microsensors on a polyp scale (a-b) and with O_2_ gas exchange measurements of whole fragments in incubation chambers (c-d). Measurements were performed on daily fed corals (a, c) and unfed corals (b, d) for each of three temperature treatments; T0 (control, solid lines), T_1_ (3 days at 30°C; dashed lines) and T_2_ (8 days at 30°C; dotted lines). Symbols with error bars represent means ± SEM (n = 12).

### Cell specific O_2_ flux for zooxanthellae

Cell specific rates of gross photosynthesis for intact corals showed an enhanced O_2_ production in fed corals compared to unfed corals at both normal and increased temperatures (Figure4). Under increased temperature, we found a significant enhancement of cell-specific photosynthetic rates, with up to 3-fold increase in O_2_ production in fed corals at T_2_ with a photon irradiance of 200 and 400 μmol photons m^-2^ s^-1^ (Figure 4a) compared to fed corals at T0 (Student’s *t*-test: *p* < 0.001; Welch’s *t*-test: *p* = 0.042, respectively). A similar trend was present for unfed corals as O_2_ production increased towards higher irradiance under heat-stress.

Freshly isolated symbionts also showed enhanced cell specific gross photosynthesis rates with feeding. They, however, showed no significant change upon increased temperature, except fed T_2_ at 50 μmol photons m^-2^ s^-1^ (Figure 4b) showing a clear enhanced O_2_ production relative to the control (T_0_) measurements (1x 10^-7^ ± 1.7x 10^-8^ *vs* 3.5x 10^-7^ ± 6.5x 10^-8^ μmol O_2_ cell^-1^ h^-1^, respectively; mean ± SEM). However, we found increased O_2_ production in fed corals compared to unfed corals for both isolated symbionts, and symbionts *in vivo* (Student’s *t*-test: *p* < 0.001; isolated symbionts from fed corals compared to unfed corals at T_0_).

### Carbon budget of entire coral fragments (macroscale)

Carbon flux was strongly influenced by both the feeding status and heat stress of *P. damicornis*. Gross production rates were enhanced under increased temperature and daily feeding (Figure 1c, d). Respiration rates (*R*_S_) of symbionts in control corals (T_0_) showed a similar rate of respired carbon for both fed and unfed corals before induced heat stress (^~^14-14.2 %; Table 1), and during heat stress (T_1-2_; 5.3-6.9 %; Table 1).

**Table 1.**
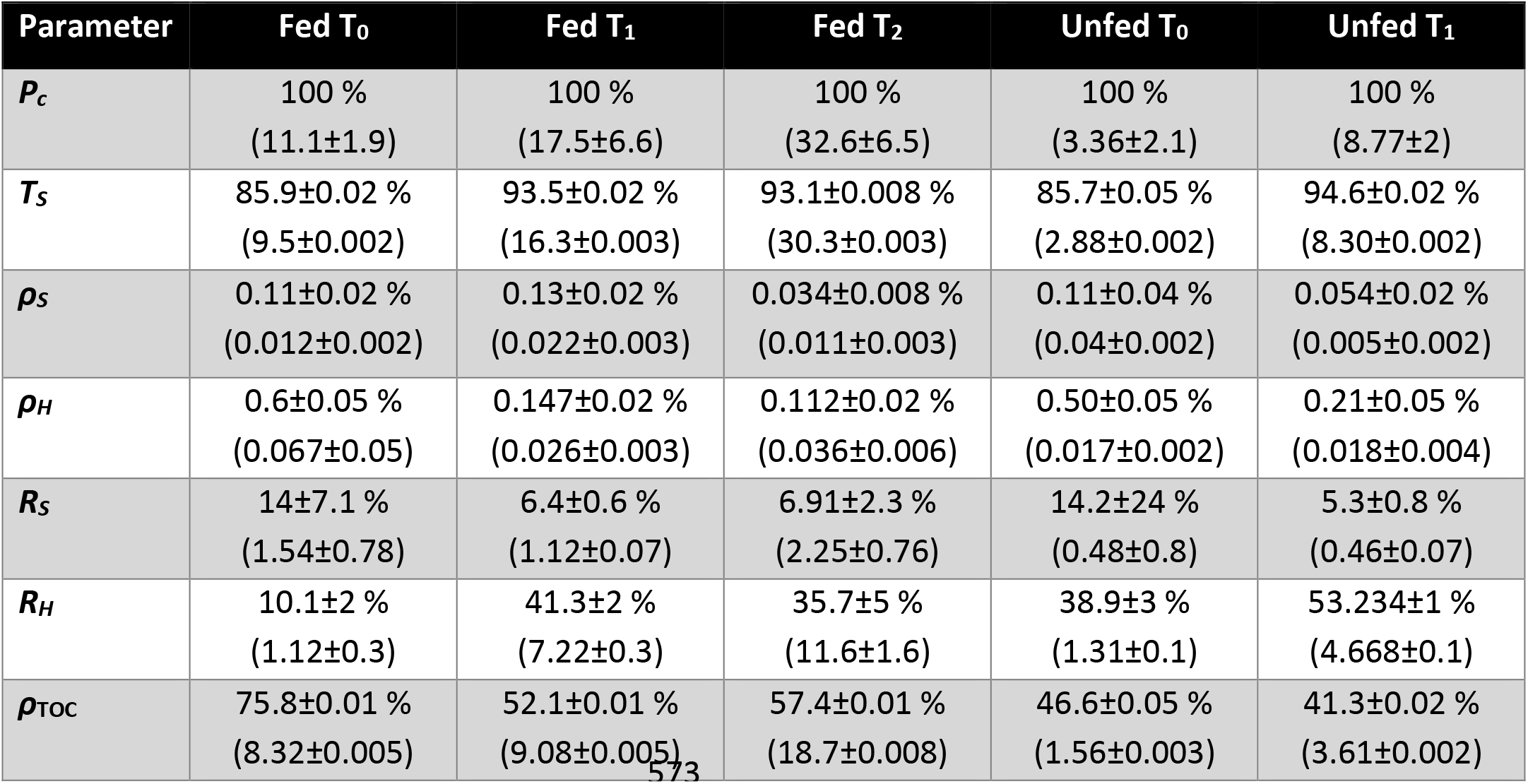
Mass balance of photosynthate translocation and carbon budget for fed and unfed *Pocillopora damicornis* as a function of time under heat stress. Incorporation rates of carbon are based on ^13^C-isotope incubation experiments using a pulse period of 4 hours. See Figure 3 for a schematic overview of the carbon budget and abbreviations used. Data represent means ± SEM (n = 3-5) and values in brackets have the unit *μg C cm*^-2^ *h*^-1^.

| Parameter | Fed T<sub>0</sub> | Fed T<sub>1</sub> | Fed T<sub>2</sub> | Unfed T<sub>0</sub> | Unfed T<sub>1</sub> |
| --- | --- | --- | --- | --- | --- |
| $P_c$ | 100 %<br>(11.1±1.9) | 100 %<br>(17.5±6.6) | 100 %<br>(32.6±6.5) | 100 %<br>(3.36±2.1) | 100 %<br>(8.77±2) |
| $T_s$ | 85.9±0.02 %<br>(9.5±0.002) | 93.5±0.02 %<br>(16.3±0.003) | 93.1±0.008 %<br>(30.3±0.003) | 85.7±0.05 %<br>(2.88±0.002) | 94.6±0.02 %<br>(8.30±0.002) |
| $\rho_s$ | 0.11±0.02 %<br>(0.012±0.002) | 0.13±0.02 %<br>(0.022±0.003) | 0.034±0.008 %<br>(0.011±0.003) | 0.11±0.04 %<br>(0.04±0.002) | 0.054±0.02 %<br>(0.005±0.002) |
| $\rho_H$ | 0.6±0.05 %<br>(0.067±0.05) | 0.147±0.02 %<br>(0.026±0.003) | 0.112±0.02 %<br>(0.036±0.006) | 0.50±0.05 %<br>(0.017±0.002) | 0.21±0.05 %<br>(0.018±0.004) |
| $R_s$ | 14±7.1 %<br>(1.54±0.78) | 6.4±0.6 %<br>(1.12±0.07) | 6.91±2.3 %<br>(2.25±0.76) | 14.2±24 %<br>(0.48±0.8) | 5.3±0.8 %<br>(0.46±0.07) |
| $R_H$ | 10.1±2 %<br>(1.12±0.3) | 41.3±2 %<br>(7.22±0.3) | 35.7±5 %<br>(11.6±1.6) | 38.9±3 %<br>(1.31±0.1) | 53.234±1 %<br>(4.668±0.1) |
| $\rho_{\text{TOC}}$ | 75.8±0.01 %<br>(8.32±0.005) | 52.1±0.01 %<br>(9.08±0.005) | 57.4±0.01 %<br>(18.7±0.008) | 46.6±0.05 %<br>(1.56±0.003) | 41.3±0.02 %<br>(3.61±0.002) |

Host respiration rates (*R*_H_ in % fixed carbon) were ^~^28.8 % higher for fed corals compared to unfed corals at T_0_ (Table 1). Heat stress enhanced host respiration for fed corals at T_1-2_ by about 1.5-fold compared to fed corals at T_0_ (Table 1; Welch’s *t*-test: *p* = 0.009; fed corals T_0→1_ and T_0→2_). Carbon translocation rates (T_s_) from symbiont to host showed a slight increase during heat stress for all treatments (85.8 ± 0.1 *vs* 93.7 ± 0.5 %; Table 1; T_1-2_ compared to T_0_). The carbon incorporation rates were low for both symbionts (*ρ_S_*) and hosts (*ρ_H_*) in all treatments as compared to other assessed coral species (Tremblay *et al*., 2016), and incorporation accounted for only 0.03-0.6 % of the fixed carbon (Table 1). A high excretion of carbon was calculated, reaching a release of between ^~^50-75 % of the initially fixed carbon in all treatments (Table 1).

## Discussion

### Implications of measuring O_2_ production at different spatial scales

The O_2_ production and consumption in *P. damicornis* were assessed with methods operating on two different spatial scales, i.e., polyp scale (mm resolution, microscale) and fragment scale (cm resolution, macroscale; Figure 1 and 2). Micro- and macro-scale measurements revealed enhanced O_2_ evolution rates for fed corals compared to unfed corals (Figure 1 and 2). However, gross O_2_ production on a macroscale was increased at high temperature (Figure 1c, d and 2c, d) compared to gross O_2_ production on a microscale, especially at T_1_ ➔ T_2_ where rates were 2.4-fold enhanced on a macroscale (Figure 1c). Our microscale measurements were restricted to the polyp region, at a defined angle near the apical tips of the coral branch, which could potentially underestimate the productivity of polyp areas that were not assessed with microsensor measurements. The difference between micro- and macroscale measurements for fed corals at T_2_ (Figure 1a, c) could thus be that photosynthesis was heterogeneously distributed over the coral surface (Al-Horani *et al*., 2005). Scalar irradiance on bare skeletons of *P. damicornis* (Supplementary figure 5) revealed that light exposure near the base of the coral branch was increased by about 100 % compared to the apical branch tip. Previous studies on spatial distribution of photosynthesis across the tissue surface revealed that the majority of photosynthetic activity is located around the tissue covering the corallite septa, with up to one order of magnitude increased photosynthesis compared to adjacent coenosarc tissue (Al-Horani *et al*., 2005). These findings emphasize the potential underestimation of the photosynthetic activity of the polyp regions near the base of the coral branch (Supplementary figure 5). The O_2_ production observed for macroscale measurements of fed corals at T_2_ may thus be additionally affected by the backscattering properties in polyp areas around the base of the coral branch, as fed corals at T_2_ had reduced cell density compared to controls (T_0_; figure 3a in Lyndby *et al*., submitted), which cause an enhanced radiative exposure of the symbionts residing in the deeper tissue layers leading to an increased O_2_ production.

Cell specific rates of O_2_ production (Figure 4a) were similar to areal estimates measured on intact coral fragments during the course of the experiment (Figure 4 vs Figure 1). Chlorophyll *a*+*c*_2_ and cell density per area decreased with increased temperature (Figure 3b in Lyndby *et al*., submitted), while chlorophyll *a*+*c*_2_ per cell values remained more or less constant (Figure 3c in Lyndby *et al*., submitted). Fewer symbionts within the coral tissue lead to reduced algal self-shading (Lesser *et al*., 2000; Terán *et al*., 2010; Wangpraseurt *et al*., 2012), increasing the light exposure per cell and thus higher cell specific photosynthetic rates (Enríquez *et al*., 2005; Wangpraseurt *et al*., 2017).

**Figure 3.**
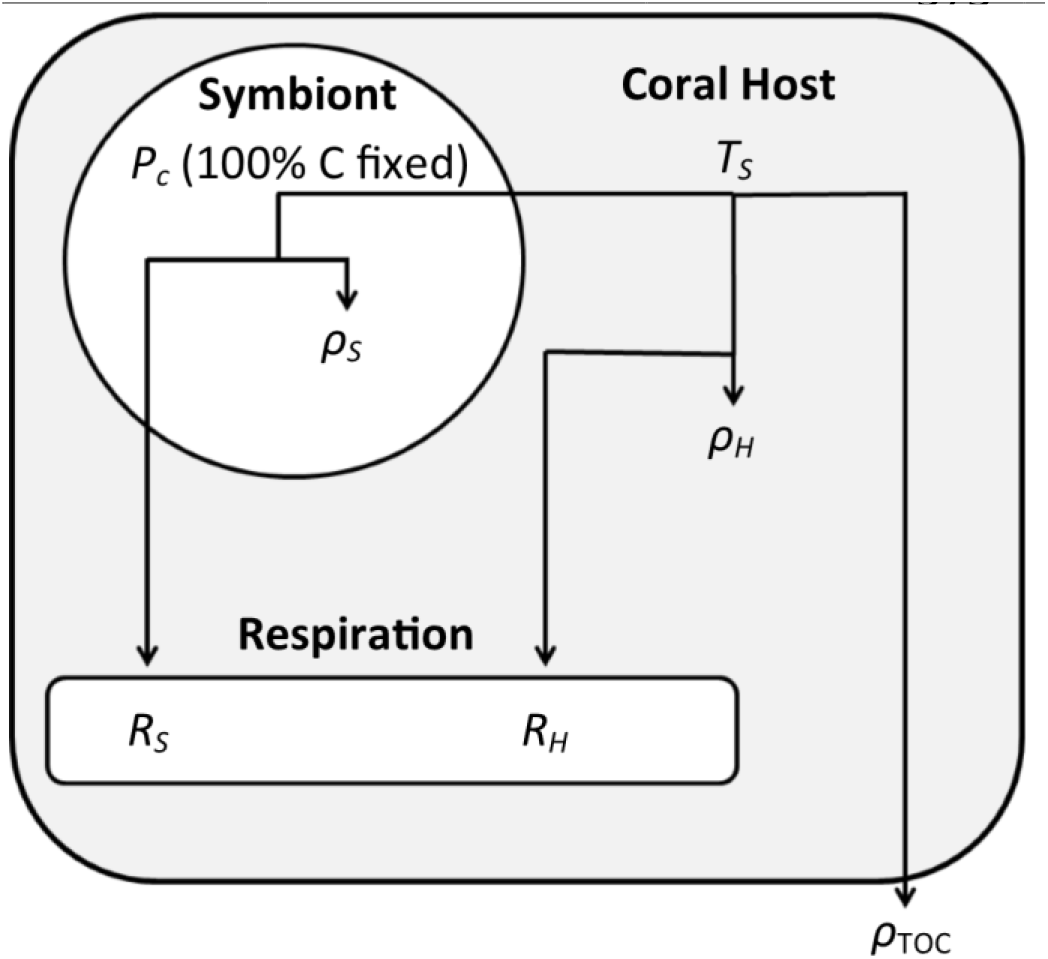
Schematic overview of photosynthate translocation and the carbon budget overall. *Pc* is the total amount of inorganic C fixed through photosynthesis, hence 100 % fixed for all treatments and time points (see Table 1 for detailed data). *T_S_* is the C translocated from symbiont to coral host, *ρ* is the C incorporated into the symbionts (*ρ_S_*) and coral host (*ρ_H_*). *R* is the C used for respiration by symbionts (*R_S_*) and by the coral host (*R_H_*). Finally, *ρ*_TOC_ is the total amount of C lost from the holobiont as organic carbon.

Gross O_2_ production rates of isolated zooxanthellae cells (Figure 4b), were about one order of magnitude lower than the measurements performed *in vivo* (Figure 4a). This suggests that the symbionts benefit from the animal tissue e.g. through optimized light exposure and potentially other host-affected factors such as pH, nutrients or spectral filtering by host pigments, leading to enhanced *in vivo* photosynthetic efficiency (Enríquez *et al*., 2005). Even though both *in vivo* and *in vitro* measurements were performed under identical levels of incident downwelling irradiance, the actual scalar irradiance will differ between the two measurements because of light scattering in the coral tissue and on the skeletal surface (Wangpraseurt *et al*., 2012, 2014). However, inorganic nutrient supply from the coral host (e.g. NH_4_^+^) can also contribute to the observed enhancement of photosynthetic efficiency *in vivo* (Borell & Bischof, 2008).

**Figure 4.**
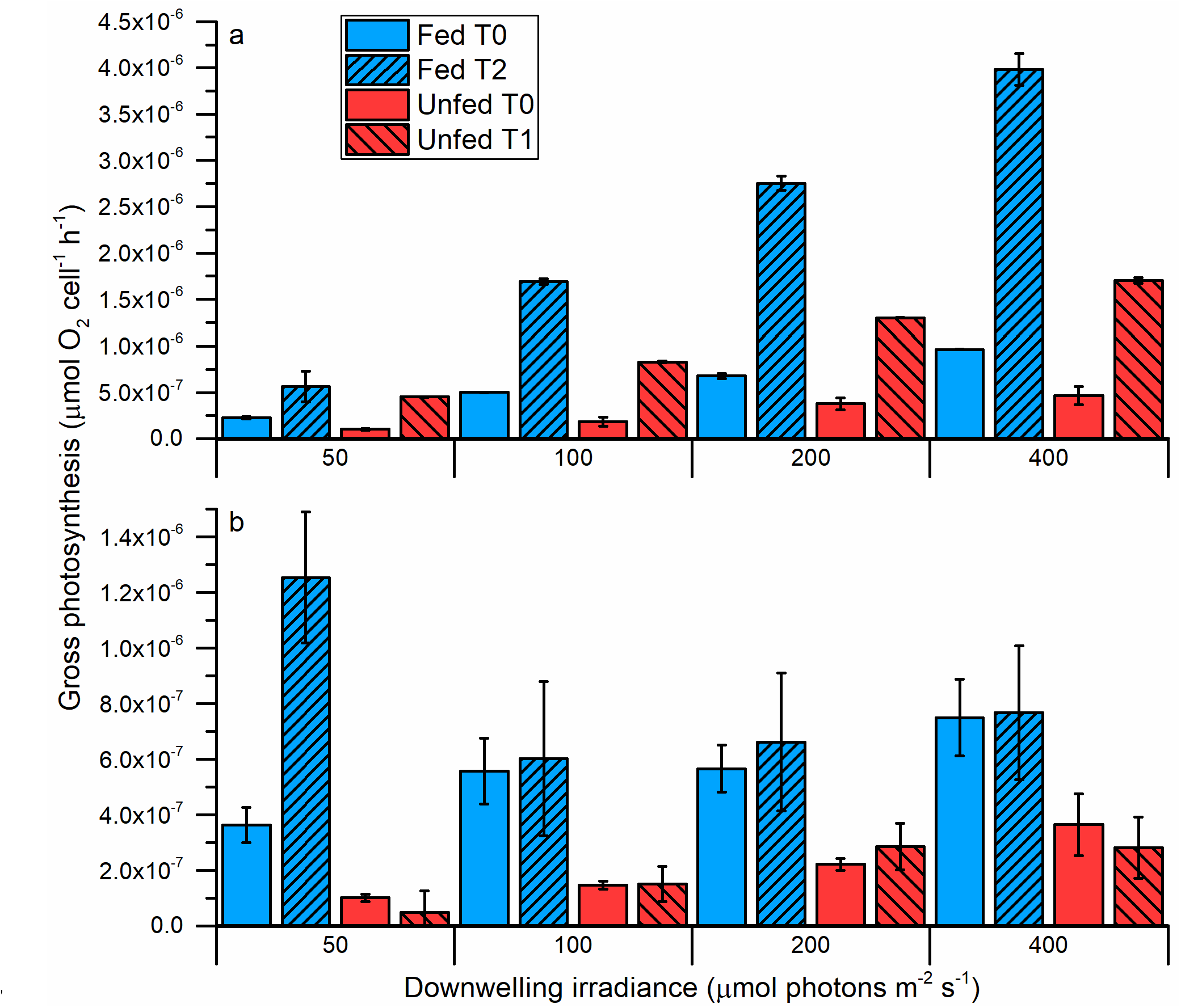
Cell specific gross photosynthetic O_2_ production rates as a function of incident photon irradiance measured on intact fragments of fed and unfed *Pocillopora damicornis* (*in vivo*, panel a) and freshly isolated zooxanthellae from *P. damicornis* (*in vitro*, panel b) at increasing exposure time to heat stress. Blue columns represent measurements on fed corals, and red columns measurements on unfed corals. Data are means ± SEM (n= 3-5).

### O_2_ dynamics during heat stress

The O_2_ dynamics of *Pocillopora damicornis* measured on a whole fragment scale (macroscale) and polyp scale (microscale) agree with previous studies from other coral species showing a 35-60% increase in chlorophyll per area and an increased cell density in fed corals (Muscatine *et al*., 1989; Fitt & Cook, 2001; Ferrier-Pagès *et al*., 2003). In contrast, the lack of host feeding in unfed corals could explain the lower cell density for both whole fragment scale and polyp scale, as sufficient nutrients to produce e.g. antioxidants to cope with ROS accumulation are not met, causing ingestion or expulsion of excess zooxanthellae (Figure 1c-d, Figure 2c-d; Figure 3 in Lyndby *et al*., submitted).

Earlier studies on thermally stressed *P. damicornis* revealed increased feeding rates of up to 127 % compared to fed controls (Courtial *et al*., 2017), implying an increased demand of nitrogen to withstand stress factors by synthesizing protective compounds (e.g. antioxidants and heat-stress proteins) to avoid oxidative stress (Downs *et al*., 2002).

Our results suggest that the nutritional input from active feeding enables the coral host to maintain a higher photosynthetic activity as compared to unfed corals, and we speculate that feeding enhances essential nutrients (e.g. nitrogen) for synthesis of antioxidants and heat-stress proteins to better prevent damage from reactive oxygen species (ROS; Demmig-Adams & Adams, 2002; Fitt *et al*., 2009).

### Autotrophic carbon fluxes and mass balanced results of photosynthates and translocation

The C incorporation rates, i.e., the rate of inorganic carbon assimilated per area and time (μg C cm^-2^ h^-1^) of the initially fixed inorganic carbon, were surprisingly low for both symbionts and the animal host in all treatments (Table 1). In contrast, previous studies found several orders of magnitude higher incorporation rates for the branching coral *Stylophora pistillata*, which belongs to the same family as *P. damicornis* (Pocilloporidae; Tremblay *et al*., 2012). The majority of autotrophically acquired carbon is used for daily household metabolism, and colonies growing in high light assimilate a very small amount of photosynthates, due to lack of nitrogen (Falkowski *et al*., 1984). This might explain the very high excretion of translocated carbon in *P. damicornis* observed in all our treatments (Table 1). A high amount of fixed carbon could surpass the metabolic demand and would therefore be excreted from the coral host, as sufficient supplies of nutrients are available from heterotrophy for fed corals (Fed T_0-2_; Table 1). Studies on energy and carbon dynamics in *Pocillopora eydouxi* (Davies, 1984) show a similar high release of photosynthates (about 50 % of fixed carbon) from coral to ambient water, which correlates well with our results of *P. damicornis* (Fed T_1-2_; Table 1).

This further supports the idea that the growth of *P. damicornis* is limited by the absence of organic nitrogen, despite the availability of autotrophic carbon (Davies, 1984). Excess carbon can also maintain a food source for surface associated microbes (Bourne & Munn, 2005; Thompson *et al*., 2015) that may confer benefits to the coral holobionts, e.g. in terms of better resilience to heat stress (Rosenberg *et al*., 2007) or nutrient supply via fixed nitrogen from diazotrophs being translocated to the coral host (Bednarz *et al*., 2017).

The host respiration for unfed corals compared to fed corals at T_0_ was 10.2 % and 38.9 % of the initially fixed carbon, respectively (Fed and unfed T_0_; Table 1). This could indicate a host stress response in unfed corals due to lack of organic nutrients, which are readily available for fed corals (Ferrier-Pagès *et al*., 2003). Additionally, studies on host response to thermal stress revealed a higher production of super oxide dismutase and heat stress proteins (Fitt *et al*., 2009), which will naturally increase the host metabolism, and correlates well with our observation of an increased host respiration in both fed and unfed corals at T_1-2_ (Table 1). We also see a clear reduction in the P:R ratio during heat stress for unfed corals from 2.56 at T_0_ down to 1.88 at T_1_ (Table 1), in line with previous assessments of thermally stressed *P. damicornis* (Coles & Jokiel, 1977).

The translocation rates of the fixed carbon going from the symbionts to the animal host seem to deviate from previously assessed corals from the Pocilloporidae family. Whereas the translocation rate usually decrease during heat stress, it remained high for *P. damicornis* throughout the treatments (Tremblay *et al*., 2012). Therefore, contrary to other coral species whose zooxanthellae keep more autotrophic carbon for their own needs and growth during thermal stress, symbionts of *P. damicornis* in this experiment always translocated more than 85 % of the acquired carbon. This may be due to the lack of other nutrients, which prevent an efficient use of the photosynthesized carbon.

## Conclusion

By applying multiple methods to determine carbon translocation and photosynthetic efficiency in fed and unfed specimens of the scleractinian coral *Pocillopora damicornis* during heat-stress, we found that carbon budgets showed low incorporation rates and high release rates of autotrophically fixed carbon compared to closely related coral species, which emphasizes the benefits of heterotrophically acquired nutrients for *P. damicornis*. Heterotrophy might thus cover a larger portion of the nutritional demand for *P. damicornis* than previously assumed, and underlines how active feeding plays a fundamental role in metabolic dynamics and bleaching susceptibility.

## Acknowledgement

We thank the staff at the Scientific Center of Monaco for their excellent assistance with experimental procedures and labor intensive general maintenance and care of the corals for the duration of this study. The study was funded by grants from the Carlsberg Foundation (DW), a Sapere Aude Advanced grant from the Independent Research Fund Denmark (MK), and the Scientific Center of Monaco (part of the RTPI Nutress).

